# NET1/ARHGEF8 is a mechanosensitive Guanine Exchange Factor controlling vascular smooth muscle cells’ contractility

**DOI:** 10.1101/2025.11.26.690669

**Authors:** Mary-Adel Mrad, Surya Batta, Virginie Vignard, Marc Rio, Gilliane Chadeuf, Eloise Gros, Vincent Sauzeau, Gervaise Loirand, Anne-Clémence Vion

## Abstract

Cyclic stretch, generated by pulsatile blood pressure, has been identified as a pivotal regulator of vascular smooth muscle cell (VSMC) behavior and arterial wall homeostasis. Research has demonstrated that RhoA activation is a fundamental component of VSMC responses to mechanical stress through its action on cytoskeletal remodeling. However, the upstream mechanosensitive regulators of RhoA activation remain insufficiently defined. In this study, we identify the guanine nucleotide exchange factor (RhoGEF) NET1/ARHGEF8 as a stretch-responsive protein. NET1 appears to be highly expressed in arteries and VSMCs. Under physiological levels of cyclic stretch, NET1 localizes to the cytosol and interacts with RhoA. In contractile cells, loss of NET1 blunted stretch-induced MYPT1 phosphorylation and impaired cell adhesion and spreading, without exerting any effect on cell proliferation. As we demonstrated, NET1’s contribution to the stretch-induced response of VSMC was due to its localization in the cytosol. The expression of a cytosolic NET1 mutant promoted contractile gene expression and increased cell contractile capacity. Collectively, these findings identify NET1 as a stretch-sensitive RhoGEF in VSMCs that contributes to their adhesion and contractile behavior.

## Introduction

In the human body, cellular behavior and tissue integrity are influenced by the surrounding environment and the associated mechanical forces. The majority of cells are indeed equipped with mechanosensors and mechanotransduction pathways, which enable them to perceive and respond to mechanical cues. This ensures proper development and maintenance of homeostasis, as well as adaptation to physiological demands. The vascular system is one of the organs that permanently experiences strong and variable physical forces. Vascular smooth muscle cells (VSMCs), located in the medial layer of the arterial wall, are continuously subjected to cyclic stretch generated by pulsatile blood flow. VSMCs can detect variations in mechanical strain and in stiffness within their surrounding extracellular matrix (ECM), allowing them to dynamically adjust their status and functions ^1–5^. Indeed, they exhibit remarkable phenotypic plasticity, existing along a spectrum from a highly contractile state (low proliferation, efficient electromechanical coupling) to a synthetic state (high ECM production, migration, secretory activity)^6^. The mechanoadaptive behavior of VSMCs is fundamental in regulating vascular tone, remodeling of vessel walls, thereby contributing to long-term blood pressure control^7,8^. Conversely, impaired mechano-responsiveness or chronic exposure to abnormal mechanical stimuli can drive inappropriate remodeling and pathological processes^8,9^, leading to conditions such as hypertension, vascular stiffening or aneurysms^10–12^.

A pivotal regulator of cell response to mechanical stimuli is the RhoA signaling. RhoA, a member of the Rho family of small G proteins, orchestrates cytoskeletal dynamics, contractility, migration, proliferation, and phenotypic switching in VSMCs^13^. Through its downstream effector Rho Kinase (ROCK), RhoA enhances calcium sensitivity of the contractile apparatus by inhibiting myosin light chain phosphatase, and regulates actin polymerization, thereby generating long-lasting contraction and vascular tone^14^. RhoA/ROCK also drives migration through focal adhesion turnover^15–17^, proliferation via cross-talk with the MAPK and PI3K signaling and also controls transcriptional programs to coordinate growth and differentiation^18^. When dysregulated, these signaling pathways favor dedifferentiation, loss of contractile markers, and adoption of a synthetic phenotype placing this axis at the heart of both vascular homeostasis and pathology^19–21^.

Upstream to RhoA, guanine nucleotide exchange factors (RhoGEFs) regulate its activation by catalyzing the exchange of GDP for GTP. In VSMCs, the three most studied RhoGEFs are LARG (ARHGEF12), which is necessary for the myogenic response in mesenteric arteries^22^, ARHGEF1 (p115-RhoGEF) and p63RhoGEF, which are both activated by angiotensin II to regulate blood pressure and proliferation, respectively^23,24^. Although VSMCs are exposed to mechanical forces, no RhoGEF has yet been clearly identified as mechanosensitive in these cells. Conversely, the literature contains a substantial amount of information on mechanosensitive RhoGEFs in other cell types^25^. Focusing on vascular cells, ARHGEF1 and ARHGEF12 respond to tensil forces in fibroblasts and endothelial cells (ECs)^26,27^, GEF-H1 responds to cyclic stretch in fibroblasts^28^, ARHGEF40 (Solo), Abr, alsin, ARHGEF10, Bcr, ARHGEF28, PLEKHG1, P-REX2, and α-PIX respond to cyclic stretch and Trio and ARHGEF18 are sensitive to shear stress in ECs ^29,30,31^. The presence of a multitude of mechanosensitive RhoGEFs in various cell types suggests that VSMCs may also possess additional mechanosensitive RhoGEFs that have not yet been identified. These unidentified RhoGEFs may play a role in VSMC contractility, phenotypic maintenance/switching, and potentially contribute to the development of vascular diseases.

Here, we identified NET1 (ARHGEF8) as a novel stretch-sensitive RhoGEF in VSMCs. We demonstrated that NET1 is a RhoA-specific RhoGEF that is translocated from the nucleus to the cytosol upon cyclic stretch and activated in contractile cells. This specific localization enhances its interaction with RhoA and contributes to the establishment of cell adhesion and cell contractile phenotype.

## Results

### Cyclic stretch regulates NET1 expression, localization, and activity in contractile cells

To characterize Rho guanine nucleotide exchange factor (RhoGEF) expression in arteries, we performed 3’RNA sequencing on cerebral and mesenteric arteries from normotensive Wistar Kyoto (WKY) rats. Among RhoGEFs, *Net1* showed the highest expression in both vascular beds (Fig. 1A and Suppl 1A). This sequencing was also performed in cerebral arteries from spontaneously hypertensive (SHR) rats in which we observed a modest increase in Net1 transcript compared with WKY cerebral arteries (3’RNA sequencing and qPCR, Fig. 1B and 1C, respectively). To assess whether this observed up-regulation of *Net1* transcript expression was related to the force exerted on the wall by high blood pressure in SHRs we compared *Net1* expression in rat aortic smooth muscle cells (rAoSMCs) from WKY rats subjected or not to different levels of cyclic stretch *in vitro*. 3ʹSRP sequencing confirmed a strong expression of *Net1* in rAoSMCs (Suppl. Fig.1B), with a trend toward increased transcript levels under stretch conditions (3ʹSRP and qPCR; Fig. 1D and 1E, respectively). At the protein level, NET1 upregulation was minimal but significant at 10% stretch, corresponding to physiological elongation of human aortic VSMCs^32^ while increasing stretching led to an important variability in NET1 expression (hAoSMCs, Suppl Fig. 1C).

**Figure 1:**
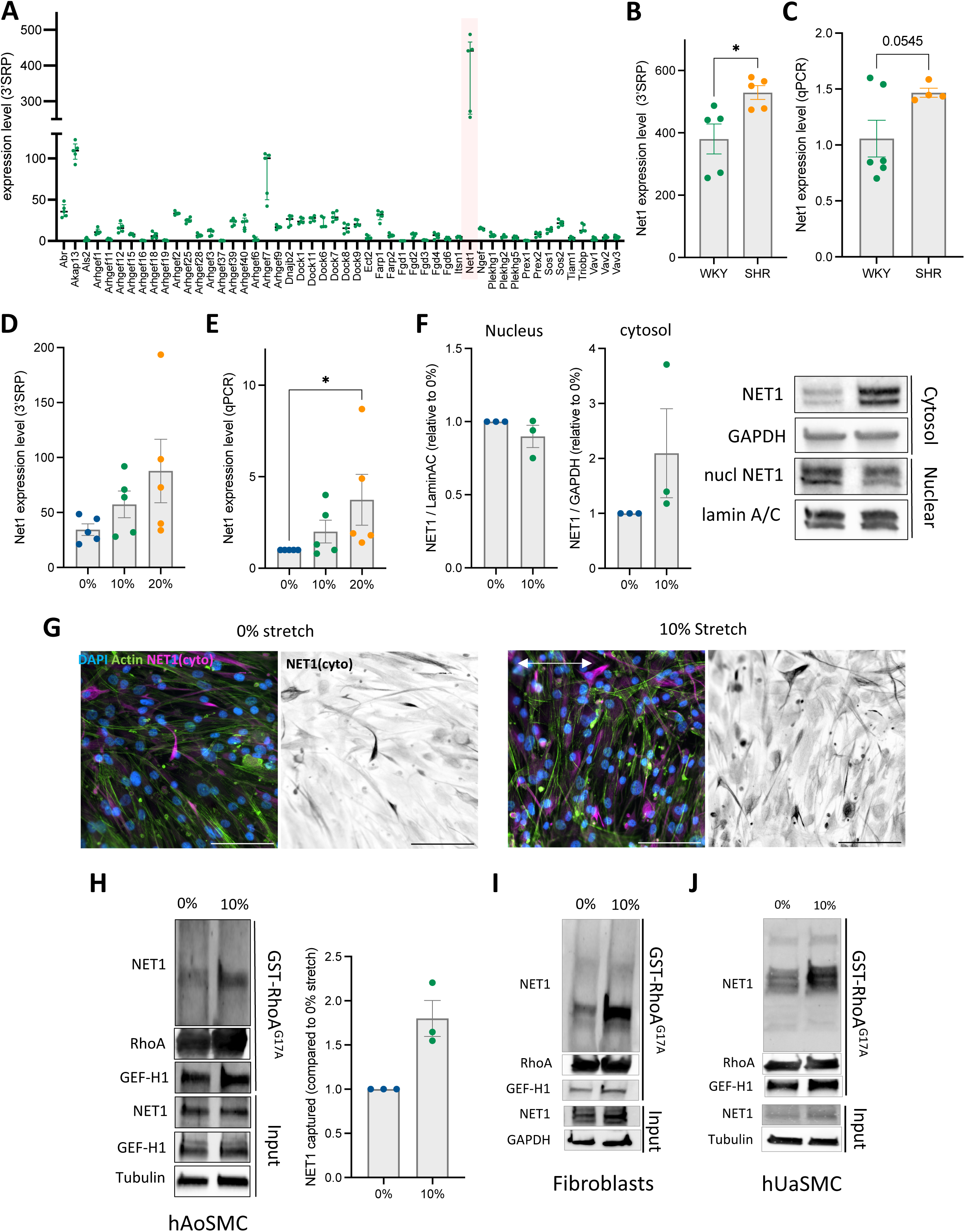
NET1 expression, localization, and activity are regulated by cyclic stretch in vascular smooth muscle cells. **(A)** In vivo quantification of RhoGEF mRNA expression by 3ʹSRP sequencing in rat cerebral arteries (N=5). Net1 expression level are highlighted in light red. **(B)** In vivo quantification of Net1 mRNA expression by 3ʹSRP sequencing in rat cerebral arteries (N=5). Unpaired T-test; *p<0.05. **(C)** In vivo quantification of Net1 mRNA expression by qPCR in cerebral arteries (N=6, WKY; N=4, SHR). Unpaired T-test. **(D)** In vitro quantification of Net1 mRNA expression by 3ʹSRP sequencing in rAoSMCs exposed to different stretch levels for 24 hours (0, 10%, 20%; 1 Hz; N=5). **(E)** In vitro quantification of Net1 mRNA expression by qPCR in rAoSMCs exposed to different stretch levels for 24 hours (0, 10%, 20%; 1 Hz; N=5). One-way ANOVA (Friedman test); *p<0.05. **(F)** In vitro quantification of nuclear and cytosolic localization of NET1 by Western blot in hAoSMCs exposed to different stretch levels for 24 hours (0, 10%; 1 Hz; N=3). GAPDH was not detected in nuclear fractions, and Lamin A/C was not detected in cytosolic fractions. **(G)** Immunofluorescent staining of NET1 in hAoSMCs exposed to different stretch levels for 24 hours (0, 10%, 20%; 1 Hz). Magenta: NET1, Green: Actin. Arrows indicate the direction of stretch. NET1 antibody from Novus (NB100-1328), recognizing human NET1 and labeling mostly the cytoplasmic NET1 on PFA fixed samples. (**H)** In vitro capture of RhoA-bound NET1 by GST-RhoA^G17A^ pull-down assay in hAoSMCs exposed to different stretch levels for 24 hours (0, 10%; 1 Hz; N=3). GEF-H1 was included as a positive control to validate the assay. **(I)** In vitro capture of RhoA-bound NET1 by GST-RhoAG17A pull-down assay in fibroblasts exposed to different stretch levels for 24 hours (0, 10%; 1 Hz). **(H)** In vitro capture of RhoA-bound NET1 by GST-RhoA^G17A^ pull-down assay in hUaSMCs exposed to different stretch levels for 24 hours (0, 10%; 1 Hz).

**Supplemental Figure 1:**
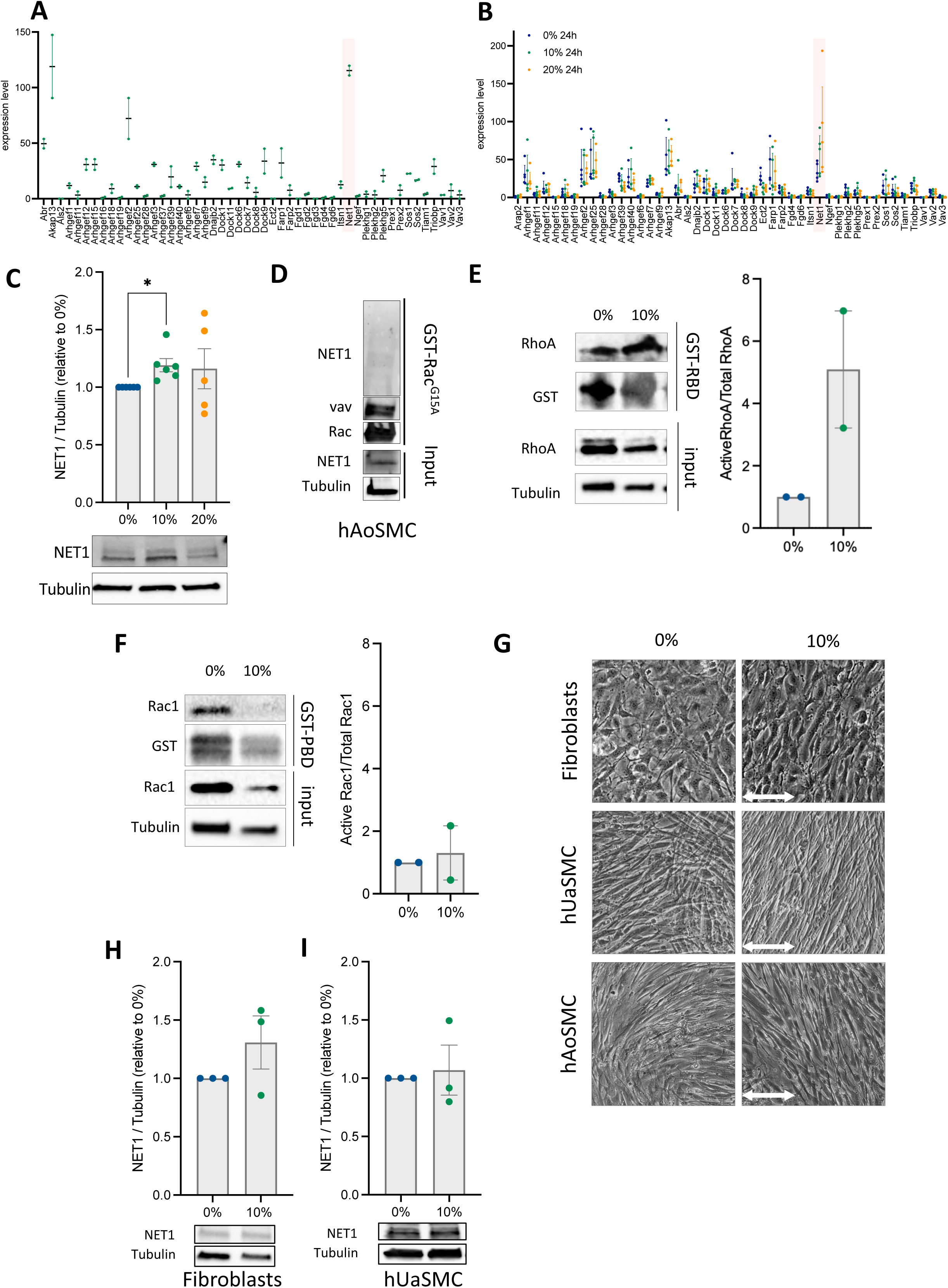
NET1 expression and activity in vascular tissues and under mechanical stretch. **(A)** 3ʹSRP sequencing data from mesenteric arteries showing expression levels of all quantified RhoGEFs (N=2). **(B)** 3ʹSRP sequencing data from rAoSMCs exposed to stretch in vitro showing expression levels of all quantified RhoGEFs (N=5). **(C)** In vitro quantification of NET1 protein expression by Western blot in hAoSMCs exposed to different stretch levels for 24 hours (0, 10%, 20%; 1 Hz; N=5). One-way ANOVA (Tukey’s post-oc); *p<0.05. **(D)** In vitro capture of Rac1-bound NET1 by GST-Rac1G15A pull-down assay in hAoSMCs. Vav was included as a positive control to validate the assay. **(E)** In vitro capture of RBD-bound RhoA by GST-RBD pull-down assay in rAoSMCs exposed to different stretch levels for 24 hours (0, 10%, 20%; 1 Hz; N=2). **(F)** In vitro capture of PBD-bound Rac by GST-PBD pull-down assay in rAoSMCs exposed to different stretch levels for 24 hours (0, 10%, 20%; 1 Hz; N=2). Y-scale is adapted to (E) to facilitate direct comparison. **(G)** Phase contrast observation of contractile cells (fibroblasts, hUaSMCs and hAoSMCs) exposed to different stretch levels for 24 hours (0, 10%; 1 Hz) representative of 3 to 5 experiments. **(H)** In vitro quantification of NET1 protein expression by Western blot in fibroblasts exposed to different stretch levels for 24 hours (0, 10%; 1 Hz; N=3). **(I)** In vitro quantification of NET1 protein expression by Western blot in hUaSMCs exposed to different stretch levels for 24 hours (0, 10%; 1 Hz; N=3).

Because NET1 contains nuclear localization signals (NLS), it can dynamically shuttle between the nucleus and cytoplasm, thereby regulating its accessibility to different downstream effectors. NET1 cytoplasmic localization allows its interaction with RhoA and activates downstream signaling while its sequestration in the nucleus prevent this binding^33^. We therefore examined whether cyclic stretch influences the subcellular distribution of NET1. In hAoSMCs, 10% stretch increased cytosolic expression of NET1 compared with unstretched controls while the nuclear pool of NET1 remain stable (Fig. 1F). This increased cytosolic localization upon stretch was also visible by immunofluorescent staining of NET1 (Fig 1G) and was accompanied by an increased NET1 binding to RhoA, as shown by pull-down assay with nucleotide-free RhoA^G17A^ mutants to capture active RhoGEF (Fig. 1H). In contrast, pull-down experiments with nucleotide-free Rac1^G15A^ mutant indicated that NET1 did not interact with Rac1, in agreement with previous reports^34^ (Suppl. Fig. 1D). These results are consistent with the increase in RhoA activity induced by 10% stretch (Suppl Fig. 1E) while Rac1 activity was not changed (Suppl Fig 1F).

Stretch-dependent regulation of NET1 was further evaluated in other cell types, the contractility of which is driven by RhoA activity including fibroblasts^35^ and human umbilical artery SMCs (hUaSMCs). When subjected to physiological stretching (10%, 1 Hz), these cells elongate and orient perpendicular to the stretch as do hAoSMCs (10% 1hz, Suppl Fig 1G). In those cells, although the total NET1 abundance upon stretch was not significantly changed (Suppl. Fig. 1H and 1I), its interaction with RhoA was drastically increased upon stretch (Fig. 1I and 1J). Together, these findings indicate NET1 is an abundant RhoGEF in the vascular wall and in VSMCs and that cyclic stretch modulates its intracellular distribution and interaction with RhoA across several types of contractile cells.

### NET1 participates in cell adhesion and contractility under stretch

To investigate the functional role of NET1 in contractile cells, we employed an shRNA-mediated silencing strategy to reduce NET1 expression in fibroblasts, which was validated by Western blot (Suppl. Fig. 2A) and immunofluorescence (Suppl. Fig. 2B). We first assessed the ability of cells lacking NET1 expression (shNET1) to adhere *in vitro*. Although the initial adhesion kinetics were not significantly different from those of control cells (shNT) (Fig. 2A, 2B), shNET1 cells displayed a reduced capacity to spread, as indicated by their lower maximum impedance (Fig. 2A, 2C). These findings suggest that NET1 contributes to cell adhesion and spreading on the extracellular matrix. When cells were subjected to mechanical stretch, one-third of the experiments done at 20% strain resulted in the detachment of shNET1 cells from their substrates (Fig. 2D), whereas no detachment was observed for shNT. In the experiments where cells remain attached, phosphorylation of Paxillin, a key regulator of focal adhesion formation and maintenance, was reduced in shNET1 cells compared to controls (Suppl Fig. 2C). Based on these observations, subsequent experiments were performed at 10% stretch to avoid cell loss and were compared to static condition. NET1 deficiency had no effect on Paxillin phosphorylation under static condition, but prevented the stretch-induced paxillin phosphorylation, confirming the specific role of NET1 in maintaining cell adhesion under stretch (Fig 2E). Similarly, phosphorylation of MYPT1, a downstream effector of RhoA/ROCK signaling, was decreased in shNET1 cells compared to shNT cells under stretch (Fig. 2F) but not under static conditions (Suppl. Fig. 2D), suggesting that NET1 is required for RhoA/ROCK-dependent cell contractility upon stretch. When observing cells morphology after stretch, NET1 deletion compromised cell orientation upon stretch, which then displayed a cell shape comparable to cell that were not exposed to stretch (Fig. 2G), comforting the idea that the mobilization of the cytoskeleton, through RhoA signaling, was impaired.

**Figure 2:**
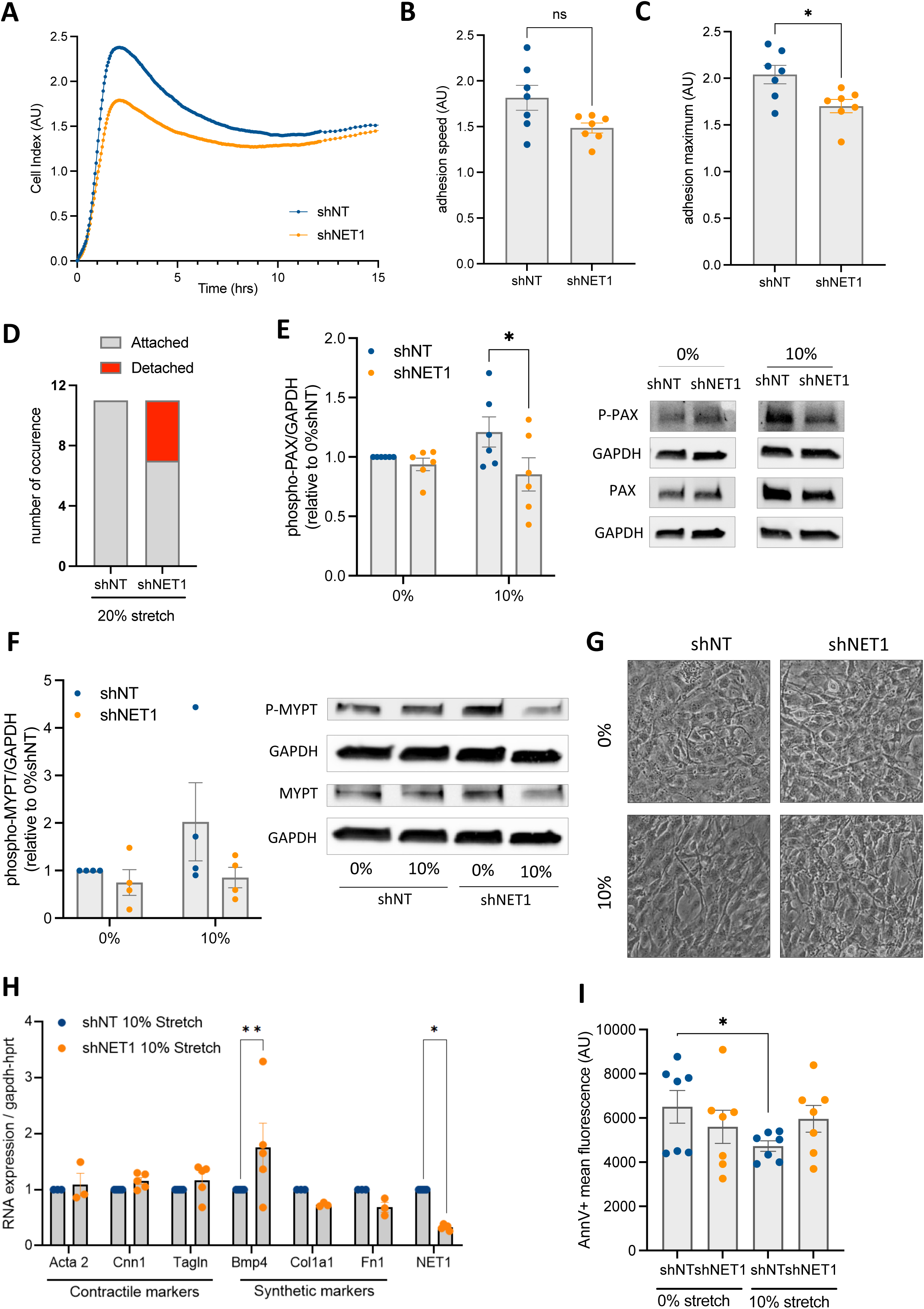
NET1 silencing modifies cell adhesion and contractility signaling but does not affect cell cycle or apoptosis under 10% stretch. **(A)** Representative adhesion curves of fibroblasts expressing shNT or shNET1 assessed by impedance under static conditions. **(B)** Quantification of the adhesion speed of fibroblasts expressing shNT or shNET1. Wilcoxon T-test. (N=7). **(C)** Quantification of the maximum of adhesion (spreading) of fibroblasts expressing a shNT or a shNET1 (N=7). Wilcoxon T-test, *p<0.05. **(D)** Quantification of the number of stretch experiments leading to cell detachment in fibroblasts expressing shNT or shNET1 (20%, 1Hz, 24 hours, 11 experiments). **(E)** Quantification and representative Western Blot of Paxillin phosphorylation in fibroblasts expressing shNT or shNET1 exposed or not to 10% stretch (1Hz, 24 hours, N=6). 2-way ANOVA, (Sidak post-oc) *p<0.05. **(F)** Quantification and representative Western blot of MYPT1 phosphorylation in fibroblasts expressing shNT or shNET1 exposed or not to 10% stretch for 24 hours (1Hz, N=4). **(G)** Phase contrast observation of fibroblasts expressing shNT or shNET1 exposed to different stretch levels for 24 hours (0, 10%; 1 Hz) representative of all experiments (<5). **(H)** Expression of contractile and secretory markers in fibroblasts expressing shNT or shNET1 exposed to 10% stretch for 24 hours (1Hz, N=3-5), evaluated by qPCR. 2-way ANOVA (Sidak post-oc), **p<0.01, *p<0.05. **(I)** Apoptosis analysis of fibroblasts expressing shNT or shNET1 to different stretch levels for 24 hours (0, 10%; 1 Hz, N=7). Annexin V mean fluorescence intensity was quantified by flow cytometry. 2-way ANOVA (Tukey’s post-oc), *p<0.05.

**Supplemental Figure 2:**
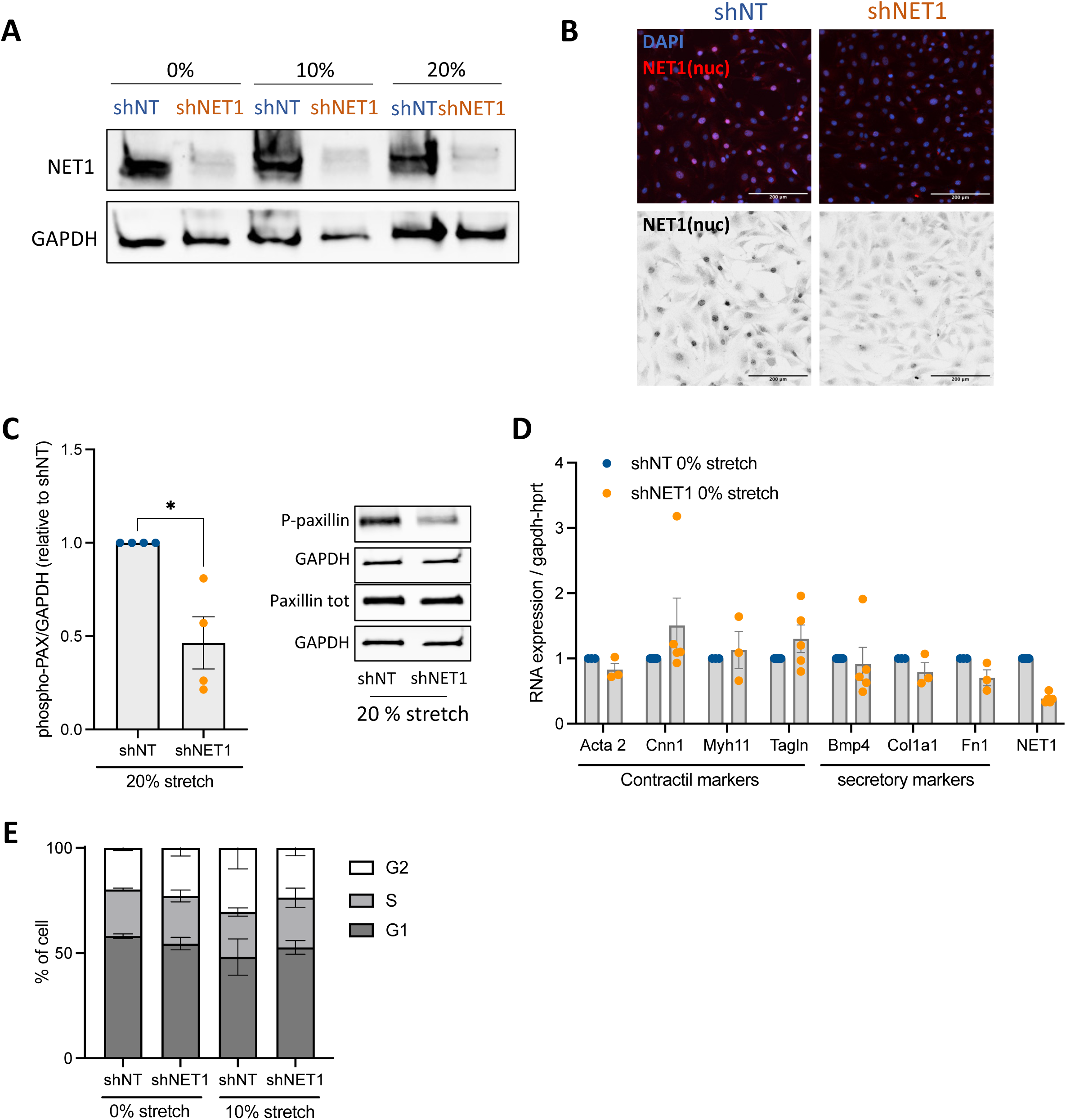
NET1 silencing modifies contractility signaling but does not affect contractile and secretory gene expression, cell proliferation, or cell cycle under static conditions. **(A)** shRNA silencing efficacy assessed by WB in fibroblasts expressing shNT or shNET1 exposed to stretch (0, 10, 20%, 24 hours, 1Hz). Representative blot of 5 experiments. **(B)** shRNA silencing efficacy assessed by immunofluorescence staining of NET1 in fibroblasts expressing shNT or shNET1. NET1 antibody from santcruz (SC50392) recognizing murin NET1 and labeling mostly the nuclear NET1. **(C)** Quantification and representative Western Blot of Paxillin phosphorylation in fibroblats expressing shNT or shNET1 exposed to stretch (20%, 1Hz, 24 hours, N=4). **(D)** Expression of contractile and secretory markers in fibroblasts expressing shNT or shNET1 under static conditions (N=3-5), evaluated by qPCR. **(E)** Cell-cycle analysis of fibroblasts expressing shNT or shNET1 under static conditions and 10% stretch (N=4), evaluated by the Two-Step Cell Cycle Assay. DAPI fluorescence histograms were used to quantify the percentage of cells in G1, S, and G2/M phases.

We next evaluated whether NET1 loss influenced the expression of contractile and secretory gene markers by qPCR. Except for *Bmp4*, which was specifically upregulated in shNET1 cells under stretch, no differences were observed in the expression of contractile or secretory genes between shNT and shNET1 cells under static conditions (Suppl. Fig. 2E) or under 10 % stretch (Fig. 2H). this suggested that NET1 was dispensable for the basal phenotypic signature of those cells both under static and stretch condition. In fibroblast, BMP4 has been described to participate in proliferation^36^. Since both stretching^37,38^ and NET1^39,40^ are also regulators of proliferation in other cell types, we investigated the effect of NET1 loss on cell proliferation. Analysis of cell cycle distribution showed no differences between shNT and shNET1 cells under either static or 10 % stretch conditions (Suppl. Fig. 2E) We then assessed the effect of NET1 loss on cell death by quantifying the phosphatidylserine exposure at the cell surface using Annexin V binding. ShNT cells exposed to stretch exhibited a lower mean fluorescence intensity of Annexin V than those under static conditions, reflecting the protective effect of physiological stretch against cell death (Fig 2J). By contrast, mean fluorescence intensity of shNET1 cells under stretch was similar to that observed under static condition suggesting that NET1 is involved in the stretch-induced signaling that protect cells against apoptosis and/or necrosis (Fig. 2J).

In summary, NET1 depletion impaired the regulation of cell adhesion, spreading, contractility and cell death induced by stretch with no effect on proliferation

### NET1 cellular localization influences VSMC function

In the previous set of experiments, NET1 was silenced in both the nuclear and cytosolic compartments, which did not allow to discriminate the relative contributions of each subcellular pool to the observed effect on stretch-induced cell response. To better understand the respective effects of the cytosolic and nuclear of NET1 on VSMCs’ functions, we generated hAoSMCs expressing either a WT-NET1, which appeared mostly sequestrated in the nucleus (Fig. 3A, left), or a NET1 mutant lacking its NLS sequences (DN-NET1), which thus localized in the cytoplasm (Fig. 3A, right), mimicking the effect of stretch in a static condition. When observing these cells, hAoSMCs expressing DN-NET1 were significantly larger than WT-NET1 cells (Fig. 3A, 3B), suggesting that cytosolic NET1 favors cell spreading. This was confirmed by impedance measurement, where DN-NET1 cells adhered faster (slope, Fig. 3C, 3D) and spread better (maximum impedance, Fig. 3C, 3E) than control and WT-NET1 cells. These results are consistent with a role of cytosolic NET1 in VSMC adhesion and spreading. We then analyzed whether the subcellular localization of NET1 could influence the phenotype of VSMCs by measuring the expression of contractile and synthetic phenotypic markers. While a complete silencing of NET1 did not affect these profiles (suppl Fig 2D), the expression of the contractile markers *Acta2* and *Cnn1* was significantly increased in DN-NET1 hAoSMCs compared to both control and WT-NET1 hAoSMCs (Fig. 3F). This increase in contractile markers translated into a functional outcome, as hAoSMC expressing the DN-NET1 retracted more a collagen gel than control and WT-NET1 cells (Fig. 3G), suggesting a stronger contractile apparatus. This finding indicates that the cytosolic localization of NET1 may promote the contractility of VSMCs.

**Figure 3:**
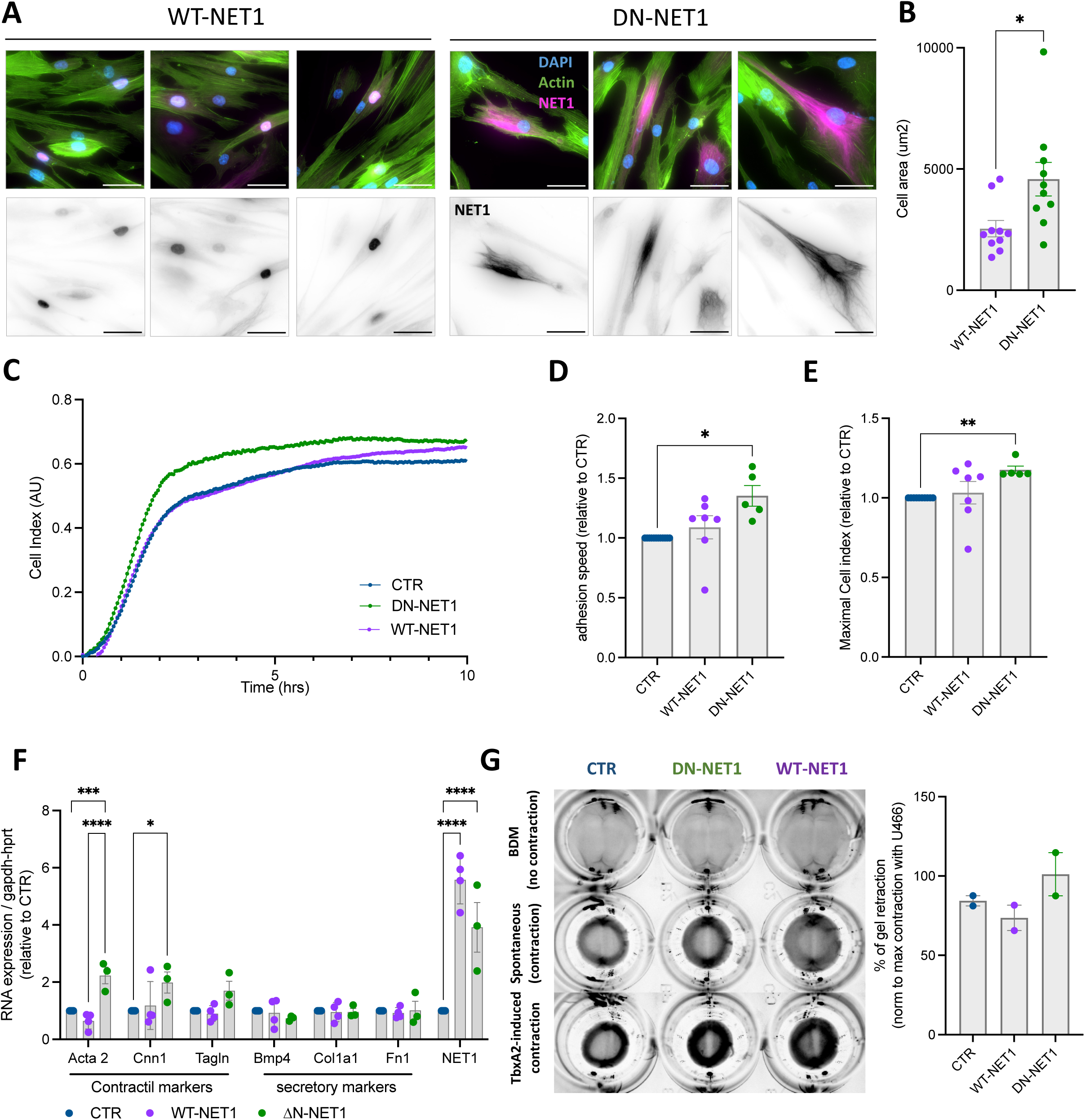
Cytoplasmic NET1 enhances cell spreading, adhesion, and contractility. **(A)** Representative images of WT-NET1 and DN-NET1 localization in hAoSMCs, assessed by immunofluorescence staining of NET1. NET1 antibody from Bethyl, recognizing human NET1 and labeling both cytosolic and nuclear overexpressed NET1. Scale bar: 10µm **(B)** Quantification of the cell area in WT-NET1 and DN-NET1 hAoSMCs (10 cells quantified from 1 infection). Unpaired T-test, *p<0.05. **(C)** Representative adhesion curves of WT-NET1 and DN-NET1 hAoSMCs, assessed by impedance, under static conditions. **(D)** Quantification of the adhesion speed of control hAoSMC or hAoSMC expressing WT-NET1 or DN-NET1 under static conditions (no stretch, N=5-7). Welsh’s T-test, *p<0.05. **(E)** Quantification of the maximum of adhesion (spreading) of control hAoSMC or hAoSMC expressing WT-NET1 or DN-NET1 under static conditions (no stretch, N=5-7). Welsh’s T-test, **p<0.01. **(F)** Expression of contractile and secretory markers in control hAoSMC or hAoSMC expressing WT-NET1 or DN-NET1, under static condition (N=3-4), evaluated by qPCR. 2-way ANOVA (Sidak post-oc), ****p < 0.0001; ***p < 0.001; *p < 0.05. **(G)** Contraction capacity of control hAoSMCs or hAoSMCs expressing WT-NET1 or DN-NET1, with or without stimulation by the thromboxane A₂ agonist (1 µM) or BDM, evaluated using a collagen gel contraction assay. Data shown and quantified at the 72-h time point (N=2).

In summary, expression of WT-NET1 and DN-NET1 suggest that the effect of NET1 on cell spreading, adhesion, differentiation and contractility are carried by cytosolic NET1, highlighting a NET1 location-dependent modulation of VSMC functions.

## Discussion

Our study identifies NET1/ARHGEF8 as a stretch-sensitive RhoGEF in contractile cells that converts cyclic mechanical strain into RhoA-dependent signaling governing cell adhesion and contractility. NET1 is a RhoA-specific GEF that resides predominantly in the nucleus under basal conditions to prevent aberrant RhoA activation at the plasma membrane^33^. Several mechanisms have been proposed to induce the export of NET1 out of the nucleus, including EGF or lysophosphatidic acid stimulation, Rac activation, JNK-, Src- or Abl1-mediated phosphorylation, or acetylation ^41,42,43^. Relocation of NET1 to the cytosol induces RhoA activation and drive RhoA-dependent signaling including actin cytoskeletal rearrangement, motility and contraction. In epithelial and cancer cells, NET1 has been shown to regulate actomyosin contractility, cell invasion ^43^, and epithelial-mesenchymal transition ^44^ through spatially restricted activation of RhoA. Here, we extend these findings to a mechanical context, demonstrating that intracellular distribution of NET1 is modulated by cyclic stretch in contractile cells, thereby suggesting that NET1 is a key component of the adaptive response of VSMCs to mechanical load.

GEF control of RhoA often relies on a change in their localization that enables their interaction with RhoA. For example, the mobility of GEFs in response to mechanical forces has been shown to be regulated by microtubule disassembly in the case of GEF-H1^45^, or by recruitment to integrin-associated complexes under tension in the case of ARHGEF12^28^. Along these lines, our data showed that NET1 appears to control RhoA activation in VSMCs upon stretching through an original relocation system involving shuffling between the nucleus and the cytosol. In our experiments, mechanical load increases the cytosolic pool of NET1, thereby facilitating close contact with RhoA during stretching. The precise origin of this accumulation in the cytosol remains unclear, as does whether it arises from new protein synthesis, altered nuclear export or cytosolic retention. As JNK and SRC have been reported to participate in the stretch response of VSMCs^46–48^, it could be hypothesized that these pathways also operate to shuffle NET1 under stretch.

In our study, NET1 silencing reduced cell spreading and adhesion and decreased paxillin phosphorylation, findings consistent with impaired focal-adhesion maturation and attenuated RhoA/ROCK-dependent signaling. These observations align with previous reports demonstrating that NET1 promotes focal-adhesion maturation through localized RhoA activation and actomyosin engagement^43^. Enforced cytosolic localization of NET1 enhanced cell adhesion and spreading, indicating that the cytosolic pool of NET1 constitutes the active fraction required for effective mechano-transduction at adhesion sites. Our data therefore suggest that NET1 could act as a mechanosensitive RhoA-GEF that couples RhoA activation to focal-adhesion assembly and tension maintenance in VSMCs under stretch.

Under physiological mechanical conditions (normal cyclic stretch and pressure, healthy ECM compliance), VSMCs maintain and reinforce their contractile phenotype^49^. The molecular basis of this process involves the RhoA-SRF(MRTF)-Myocardin transcriptional regulation axis^37,50^. As MRTFs are sequestered in the cytoplasm by G-actin binding, the shift towards F-actin by RhoA activation frees MRTFs to translocate into the nucleus, driving contractile gene expression. The precise mechanisms by which this axis is controlled in an integrated environment, including cues from pressure, matrix stifness and chemical factors, especially during vascular diseases progression, remain to be elucidated. It is interesting to note that rat hypertensive models present an increased expression of SRF and myocardin, which is responsible for the hypercontractile phenotype of VSMCs^51–53^. In the present study, enforced cytoplasmic localization of NET1 increased the expression of contractile markers in vitro. In Vivo, our data on arteries from hypertensive rats revealed an increased NET1 expression, indicating NET1’s potential role in the adaptive remodeling response to sustained mechanical stress. While the current data do not establish a link with the SRF pathway, NET1 could be a contributing factor in the modulation of VSMCs’ contractile phenotype by mechanical overload during hypertension.

In summary, NET1 emerges as a mechanosensitive RhoGEF activated by cyclic stretch, whose cytoplasmic localization is central to the regulation of VSMC adhesion, spreading, and contractility.

## Material and Methods

### Experimental model and subject details

#### Rat strains

Male Wistar Kyoto (WKY) and Spontaneously Hypertensive Rats (SHR) were purchased from Charles River and maintained under standard husbandry conditions. No experimental interventions were performed before sacrifice. Animals were euthanized with Euthasol (Dechra, pentobarbital, 140 mg/kg), and the aorta, cerebral arteries, and mesenteric arteries were collected for subsequent analyses. All the procedures were approved by the Animal Ethics Committee of Pays de la Loire (CEEA number 6). Animal care and housing were provided by the accredited animal facility at Nantes University’s UTE (Unité de Thérapeutique Expérimentale; authorization number UTE D44278).

#### Mammalian cell lines

Mouse 3T3-L1 fibroblasts, kindly provided by Dr. J. Pairault, were cultured in Dulbecco’s Modified Eagle Medium (DMEM, Thermo Fisher) supplemented with 10% fetal bovine serum (FBS), 1% L-glutamine, and 1% penicillin-streptomycin. 3T3-L1 is a sub clonal cell line derived from the original 3T3 Swiss albino cell line of 1962. Without any specific stimulation, those cells retain their fibroblast properties as the original 3T3. Human aortic smooth muscle cells (hAoSMCs) were purchased from PromoCell, derived from healthy non-hypertensive donors, and cultured in Smooth Muscle Cell Basal Medium (PromoCell) supplemented with 10% FBS, 1% penicillin-streptomycin, and corresponding supplements. Human umbilical smooth muscle cells (hUaSMCs) were purchased from Cell Systems, provided as a vial from pooled donors, and maintained in Complete Classic Cell Culture Medium (Cell Systems) supplemented with 10% FBS, 1% penicillin-streptomycin, and corresponding supplements. All cell types were cultured at 37°C in a humidified atmosphere with 5% CO₂. For functional studies, hAoSMCs and hUaSMCs were used between passages 2 and 5, before loss of their contractile phenotype and responsiveness to stretch, while 3T3-L1 fibroblasts were used between passages 15 and 20.

### Method details

#### Isolation and culture of primary rat aortic smooth muscle cells

Primary rat aortic smooth muscle cells (rAoSMCs) were isolated from thoracic aortae of male Wistar rats (8-10 weeks old). The thoracic aorta was carefully excised under sterile conditions, transferred to ice-cold phosphate-buffered saline (PBS; pH 7.4), and cleaned of surrounding connective tissue and fat under a binocular stereomicroscope. After removal of the endothelial layer, the tissue was cut into 1-2 mm fragments and subjected to enzymatic digestion with collagenase type II (2 mg/mL) prepared in high-glucose DMEM supplemented with 1% penicillin-streptomycin and 1% L-glutamine. Digestion was carried out at 37 °C for 2 h with gentle agitation, after which the reaction was stopped by adding an equal volume of DMEM containing 10% FBS. The suspension was centrifuged at 400 g for 5 min, washed twice with culture medium, and the resulting explants were seeded into culture wells containing a minimal volume of DMEM to ensure tissue-surface contact. Cells were maintained in DMEM supplemented with 10% FBS, 1% penicillin-streptomycin, and amphotericin B (10 μM) at 37 °C in a humidified 5% CO₂ atmosphere. The culture medium was replaced every 48 h until confluence. Primary cultures were subsequently expanded, and cells from passages 2 to 4 were used for the experiments.

#### In vitro mechanical stretch

Rat aortic smooth muscle cells (rAoSMCs), human aortic smooth muscle cells (hAoSMCs), human umbilical artery smooth muscle cells (hUaSMCs), and 3T3-L1 fibroblasts were subjected to uniaxial mechanical stretch using the STREX Automated Cell Stretching System (ST-1400, Greenleaf Scientific). Cells were seeded at confluence onto fibronectin-coated (2 µM) polydimethylsiloxane (PDMS) stretch chambers to ensure uniform adhesion. Stretch chambers were placed in the stretching unit and maintained within a humidified CO₂ incubator. Cyclic uniaxial stretch was applied at 10% or 20% strain for 24 h at 60 cycles per minute. After mechanical loading, cells were harvested for molecular assays (Western blotting, quantitative RT-qPCR, subcellular fractionation, flow cytometry) or fixed for morphological analyses (immunofluorescence microscopy, proliferation assays). Only experiments with control cells aligning perpendicular to the axis of strain at 10% were kept.

#### 3’Sequencing RNA Profiling

##### Experimental protocol

RNA sequencing libraries were prepared following the 3ʹ-digital gene expression (3ʹ-DGE) protocol developed by the Broad Institute (https://dx.doi.org/10.21203/rs.3.pex-1336/v1). For each sample, 10 ng of total RNA in 4 µL served as the input material. During template-switching reverse transcription, the poly(A) tails of mRNAs were tagged with universal adapters, well-specific barcodes, and unique molecular identifiers (UMIs). Barcoded cDNAs from multiple samples were subsequently pooled, amplified, and subjected to a transposon-mediated fragmentation (tagmentation) approach that enriches for 3ʹ ends of transcripts. Specifically, 200 ng of full-length cDNA were processed using the Nextera™ DNA Flex Library Preparation Kit (Illumina, #20018704) in combination with Nextera™ DNA CD Indexes (24 Indexes, 96 Samples; Illumina, #20018707), strictly following the manufacturer’s protocol (Nextera DNA Flex Library Document, #1000000025416 v04, Illumina). The resulting libraries were assessed for fragment size distribution using an Agilent 2200 TapeStation System. Libraries ranging from 350-800 bp were selected and sequenced on an Illumina HiSeq 2500 platform using the HiSeq Rapid SBS Kit v2 (50 cycles; #FC-402-4022).

##### Bioinformatics Protocol

Sequencing output consisted of paired-end fastq files. The first read contained 16 bases, comprising a 6-base sample-specific barcode and a 10-base UMI. The second read (104 bases) corresponded to the captured poly(A)-tailed RNA sequence. Bioinformatic analyses were conducted using a Snakemake workflow (https://bio.tools/3SRP). Demultiplexing of barcoded samples was performed with a custom Python script. Raw paired-end fastq files were converted into single-end fastq files for each sample. Alignment to the RefSeq reference transcriptome (downloaded from the UCSC Genome Browser) was carried out using BWA. Following alignment, UMIs were parsed and counted per gene to generate an expression matrix reflecting the absolute mRNA abundance across samples. Reads mapping ambiguously to multiple genes or exhibiting more than three mismatches relative to the reference sequence were excluded from further analysis. The final expression matrix was normalized, and differential gene expression analysis was performed using the R package DESeq2.

#### Lentiviral vector design, production, and inducible gene expression

##### Generation of Lentiviral vectors

For RNA interference, LT3GEPIR-Nontarg was constructed by amplifying the miRNA sequence having no similarity or complementary to either Human or Mouse transcriptomes and cloned in between XhoI and EcorI sites of LT3GEPIR as described by Fellmann and collaborators ^54^. The sense and anti-sense sequences were taken from Lee and collaborators ^55^. LT3GEPIR-MuNet1.1 was constructed by amplifying miRNA targeted to Mouse *Net1* and cloned between XhoI and EcorI sites of LT3GEPIR.

For NET1 overexpression constructs, the LT3HAGPIR vector was generated by amplifying GFP from pEGFPC1 using primers HAFP-Fw (5ʹ-ATTACAGATCTCGAGCCACCATGTACCCATACGATGTTCCAGATTACGCTGGATCCATGGTGAGCAAGGGCGAGGAG-3ʹ) and FP-Rev (5ʹ-GATCTGAATTCTTACTTGTACAGCTCGTC CATGC-3ʹ), which introduced an HA tag and a BamHI restriction site at the N-terminus. The amplified fragment was digested with BglII and EcoRI and cloned into the LT3GEPIR vector (Addgene #111177) to yield LT3HAGPIR. This cloning step disrupted the original BamHI site in LT3GEPIR but introduced a new BamHI site downstream of the HA tag. Full-length and N-terminally truncated Net1 fragments (MuNet1WT and MuNet1ΔN) were excised from pEF-HANet1 and pEF-HANet1-ΔN vectors, respectively, and subcloned into LT3HAGPIR between the BamHI and EcoRI sites. The resulting constructs, LT3HAGPIR-MuNet1WT and LT3HAGPIR-MuNet1ΔN, encode HA-tagged versions of NET1 under the control of the doxycycline-inducible TRE3G promoter. These onstruct also express shNRA targeting the endogenous NET1.

##### Lentivirus Production, Transduction, and Induction

Lentiviral particles were produced in HEK293T cells via transient transfection. Lentiviral transfer vectors (containing shRNA or NET1 ORF) were co-transfected with psPAX2 (packaging plasmid) and pVSVG2 (envelope plasmid) using polyethylenimine (PEI). After overnight incubation, the transfection medium was replaced with fresh DMEM supplemented with 10% fetal bovine serum and 1% penicillin/streptomycin. Viral supernatants were collected at 48 h and 72 h post-transfection, filtered through a 0.22 μm membrane, and directly applied to hAoSMCs at passage 3 and 3T3-L1 cells at passage 15 in the presence of polybrene (8 μg/mL) to enhance transduction efficiency. Following infection, cells were subjected to puromycin selection (2 μg/mL for 48 h), and surviving populations were maintained with 1 μg/mL puromycin. To induce the expression of either shRNA or NET1 constructs, doxycycline (1 μg/mL) was added at least two days before the experiment and maintained throughout the experimental procedure to sustain expression levels.

#### RT-qPCR

##### Extraction, purification, and quantification of total RNA

Cell lysis and RNA extraction were performed using the NucleoSpin RNA Plus Kit (Macherey-Nagel, 740984), following the manufacturer’s instructions. RNA concentration and purity were determined using a Nanodrop spectrophotometer (Thermo Fisher Scientific) by measuring A260/A280 and A260/A230 ratios.

##### RT-qPCR

1 µg of total RNA was used as input for complementary DNA (cDNA) synthesis using the High-Capacity cDNA Reverse Transcription Kit (Thermo Fisher Scientific), following the manufacturer’s instructions. The reverse transcription reaction was assembled with the kit’s reaction buffer, dNTPs, random primers, RNase inhibitor, and the MultiScribe™ Reverse Transcriptase, and was carried out in a total volume of 20 µL. cDNAs were stored at −20 °C until use. Quantitative real-time PCR was performed using TaqMan probe-based chemistry and the TaqMan Universal PCR Master Mix (Thermo Fisher Scientific), following the manufacturer’s recommendations. Each qPCR was carried out in a final volume of 10 µL, consisting of the Universal PCR Master Mix, 0.5 µL of the gene-specific TaqMan primer/probe mix, and diluted cDNA.

Amplifications were performed on a real-time PCR system compatible with TaqMan technology under standard cycling conditions. Data acquisition and CT determination were performed using the instrument’s analysis software. Gene expression values were normalized to the endogenous housekeeping genes 18S for rat samples and HPRT1 and GAPDH for mouse and human samples. Relative expression levels were calculated using the ΔΔCt method. The TaqMan assays used for each target gene are listed in the key resources table.

#### Subcellular fractionation analysis

Cytoplasmic and nuclear fractions were isolated using a modified Abcam protocol (Subcellular Fractionation Protocol, June 2016). All procedures were carried out at 4 °C with samples maintained on ice. Briefly, hAoSMCs grown on PDMS stretch chambers in smooth muscle cell basal medium (Promocell) were harvested and resuspended in fractionation buffer (20 mM HEPES, pH 7.4; 10 mM KCl; 2 mM MgCl₂; 1 mM EDTA; 1 mM EGTA; supplemented with 1 mM DTT and protease inhibitor cocktail). Following a 15-minute incubation on ice, cells were mechanically disrupted by repeated passage through a 27-gauge needle. Lysates were incubated for an additional 20 minutes on ice and then centrifuged at 720 × g for 5 minutes. The resulting pellet, enriched in nuclei, was washed once with fractionation buffer, further dispersed by passage through a 25-gauge needle, and re-centrifuged at 720 × g for 10 min. The purified nuclear pellet was resuspended in RIPA buffer (50 mM Tris-HCl, pH 7.5, 150 mM NaCl, 20 mM EDTA, 0.5% sodium deoxycholate, 1% NP-40, 0.1% SDS; supplemented with protease and phosphatase inhibitors) and briefly sonicated to shear genomic DNA. The supernatant from the initial centrifugation was collected as the cytoplasmic fraction. Purity of cytoplasmic and nuclear extracts was confirmed by immunoblotting against compartment-specific marker proteins, Lamin A/C (nuclear marker) and GAPDH (cytoplasmic marker).

#### Pulldown assay

Pulldown assays were performed to detect active form of Rho proteins (RhoA and Rac) and active NET1, as previously described ^56^. hAoSMCs, hUASMCs, and 3T3-L1 cells were subjected to either 10% cyclic stretch or maintained under static conditions for 24 hours. Cells were then lysed in RBD lysis buffer (50 mM Tris-HCl, pH 7.6; 500 mM NaCl; 1% Triton X-100; 0.1% SDS; 0.5% deoxycholate; 10 mM MgCl₂) for Rho protein pulldown, or GEF lysis buffer (20 mM HEPES, pH 7.5; 150 mM NaCl; 5 mM MgCl₂; 1% Triton X-100; 1 mM DTT) for NET1 pulldown, both supplemented with protease and phosphatase inhibitors. Lysates were cleared by centrifugation at 11,000 × g for 10 minutes. Equal amounts of protein (500 µg) were incubated with 50 µg of either the Rho binding domaine (RBD) of Rhotekin (GST-Rhotekin-RBD) or PAK1 (GST-PAK1-RBD) to pull-down active RhoA or Rac, or GST-RHOA^G17A^ or GST-RAC1^G15A^ to pull-down active RhoA-GEF or Rac1-GEF bound to glutathione magnetic agarose beads (Sigma) for 45 minutes at 4 °C. Beads were washed three times with the respective wash buffers, and bound proteins were subsequently detected by Western blot analysis.

#### Impedance-based assay

Cell adhesion was assessed using the xCELLigence Real-Time Cell Analysis (RTCA) system (Agilent Technologies). E-Plate 16 wells (5469830001; Agilent) were coated with fibronectin at a concentration of 2 µg/mL in PBS before cell seeding. Briefly, 3T3-L1 cells expressing either non-targeting shRNA (shNT) or shNET1.1, as well as hAoSMCs overexpressing WT NET1 or ΔN-NET1, were seeded at a density of 10,000 cells per well. Cell adhesion was monitored in real time by measuring the electrical impedance between microelectrodes integrated into the bottom of the wells. Changes in impedance were recorded and expressed as the cell index, reflecting cell attachment and spreading over the measurement period.

#### Western blotting

To assess protein expression, hAoSMCs, 3T3-L1 cells, and hUASMCs were washed with cold PBS and lysed in RIPA buffer (as described above) supplemented with protease and phosphatase inhibitors. Lysates were cleared by centrifugation, and protein concentrations were determined using the DC Protein Assay (Bio-Rad). Equal amounts of protein were mixed with Laemmli buffer (composition), denatured, and separated by SDS-PAGE using precast gels (Bio-Rad). Proteins were transferred onto nitrocellulose membranes, and equal loading was confirmed by Ponceau S staining. Membranes were blocked in 5% milk and incubated overnight with the indicated primary antibodies <u>(key resources</u> <u>table)</u>, followed by incubation with appropriate secondary antibodies (1:1000). Detection was performed using ECL (Bio-Rad), and images were captured with the ChemiDoc imaging system (Bio-Rad). Membranes were subsequently reprobed with anti-GAPDH or anti-tubulin antibodies. Protein bands were quantified using Fiji and expressed relative to GAPDH, tubulin, or as the ratio of phosphorylated to total protein.

#### In vitro Immunofluorescence staining

Immunofluorescence staining was performed in both hAoSMCs and 3T3-L1 cells to assess NET1 expression under stretch conditions and to validate the efficiency of NET1 KD, respectively. Cells were fixed with 4% paraformaldehyde (PFA) in PBS for 10 min at room temperature. Fixation was followed by blocking and permeabilization using Claudio’s blocking buffer (CBB; Franco et al., 2015) ^57^, containing 1% FBS (Gibco), 3% BSA (Sigma-Aldrich), 0.5% Triton X-100 (Sigma-Aldrich), 0.01% sodium deoxycholate (Sigma-Aldrich), and 0.02% sodium azide (Sigma-Aldrich) in PBS, pH 7.4, for 1-2 h with gentle rocking at 4 °C. Primary antibody against NET1 (Bethyl for WT-NET1/DN-NET1; Novus for human cells or SantaCruz for murin cells, 1:100) was diluted in a 1:1 mixture of CBB and PBS and incubated at room temperature for 2 h. Following washes, cells were incubated with secondary antibodies (donkey anti-rabbit, Life Technologies, 1:500) in the same buffer for 1 h at room temperature. Nuclei were labeled with DAPI (Sigma-Aldrich, 1:1000, 5 min), and polymerized actin was visualized using phalloidin-Alexa Fluor 488 (Life Technologies, 1:500, 45 min).

#### EdU Proliferation Assay

3T3-L1 cells were treated with doxycycline (1 µg/mL) for 4 days to induce plasmid-mediated NET1 silencing, and then plated in ibidi plates and allowed to adhere overnight. Maintaining doxycycline treatment, cells were incubated with EdU (10 µM) for 4 hours under static conditions (no stretch) at 37 °C and 5% CO₂. Following incubation, cells were fixed with 4% paraformaldehyde for 10 min at room temperature, washed, and permeabilized with Claudio’s blocking buffer for 30 min. Incorporated EdU was detected using a click chemistry reaction with a fluorescent azide, according to the manufacturer’s instructions (Abcam). Nuclei were counterstained with DAPI for 10 min. Images were acquired using an inverted Nikon microscope at 20× magnification. The proliferation rate was calculated as the percentage of EdU-positive cells relative to total nuclei (EdU⁺/total).

#### Cell cycle analysis

Cell cycle distribution was assessed using the NucleoCounter® NC-3000™ (ChemoMetec) following the two-step cell cycle assay protocol. Briefly, cells were harvested, centrifuged for 5 min at 400 × g at room temperature, and washed with PBS. Cells were then permeabilized and stained with DAPI (10 μg/mL) using the NucleoCounter® reagent, according to the manufacturer’s instructions. Approximately 10 µL of the stained cell suspension was loaded into the chambers of the NC-Slide A8™ (ChemoMetec). The slides were analyzed using the NucleoCounter® NC-3000™ system. The resulting histograms were analyzed using NucleoView™ software, and cells were categorized into G1, S, and G2/M phases based on their DNA content. Data are presented as the percentage of cells in each phase relative to the total cell population.

#### Apoptosis analysis by flow cytometry

Apoptosis was assessed using the Dead Cell Apoptosis Kit (Thermo Fisher Scientific), following the manufacturer’s instructions. Briefly, cells were harvested and washed with Annexin V Binding Buffer. Approximately 1 × 10⁵ cells were resuspended in 100 µL of Annexin V Binding Buffer. Five microliters of Annexin V APC were added to the cell suspension. After a 15-minute incubation at 37 °C in the dark, 400 µL of Annexin V Binding Buffer was added to each sample. Stained cells were analyzed immediately by flow cytometry using the appropriate filters for APC detection. Data were analyzed using flow-Jo to determine the percentage of apoptotic cells (Annexin V-positive) and the mean fluorescence intensity (MFI) of Annexin V.

#### Collagen gel contraction assay

Cell contraction was assessed using the Collagen-based Cell Contraction Assay Kit (Cell Biolabs, San Diego, CA) according to the manufacturer’s instructions. Briefly, cells were embedded in a collagen matrix and plated in 24-well plates. After gel polymerization, gels were released from the well surface to allow contraction. Contraction was evaluated under three conditions: spontaneous contraction (no treatment), in the presence of the myosin ATPase inhibitor BDM (at the concentration recommended by the manufacturer), and after stimulation with the thromboxane A₂ analogue U-46619 (1 µM, Santa Cruz Biotechnology). The extent of gel contraction was quantified 72 hours post-treatment by measuring the gel area using ImageJ software, and results are expressed as the percentage of area reduction relative to the initial gel area.

#### Statistical analysis

Statistical analysis was performed using GraphPad Prism software. For *in vitro* and *in vivo* experiments, when only 2 conditions were compared, a paired T-test or a Wilcoxon T-test was used, depending on the distribution of the data. For multiple comparisons, an ordinary one-way ANOVA (data distribution was assumed to be normal) was used, followed by a Tukey test. Details of the statistical test used for each experiment can be found in the figure legends.

## Key resources table

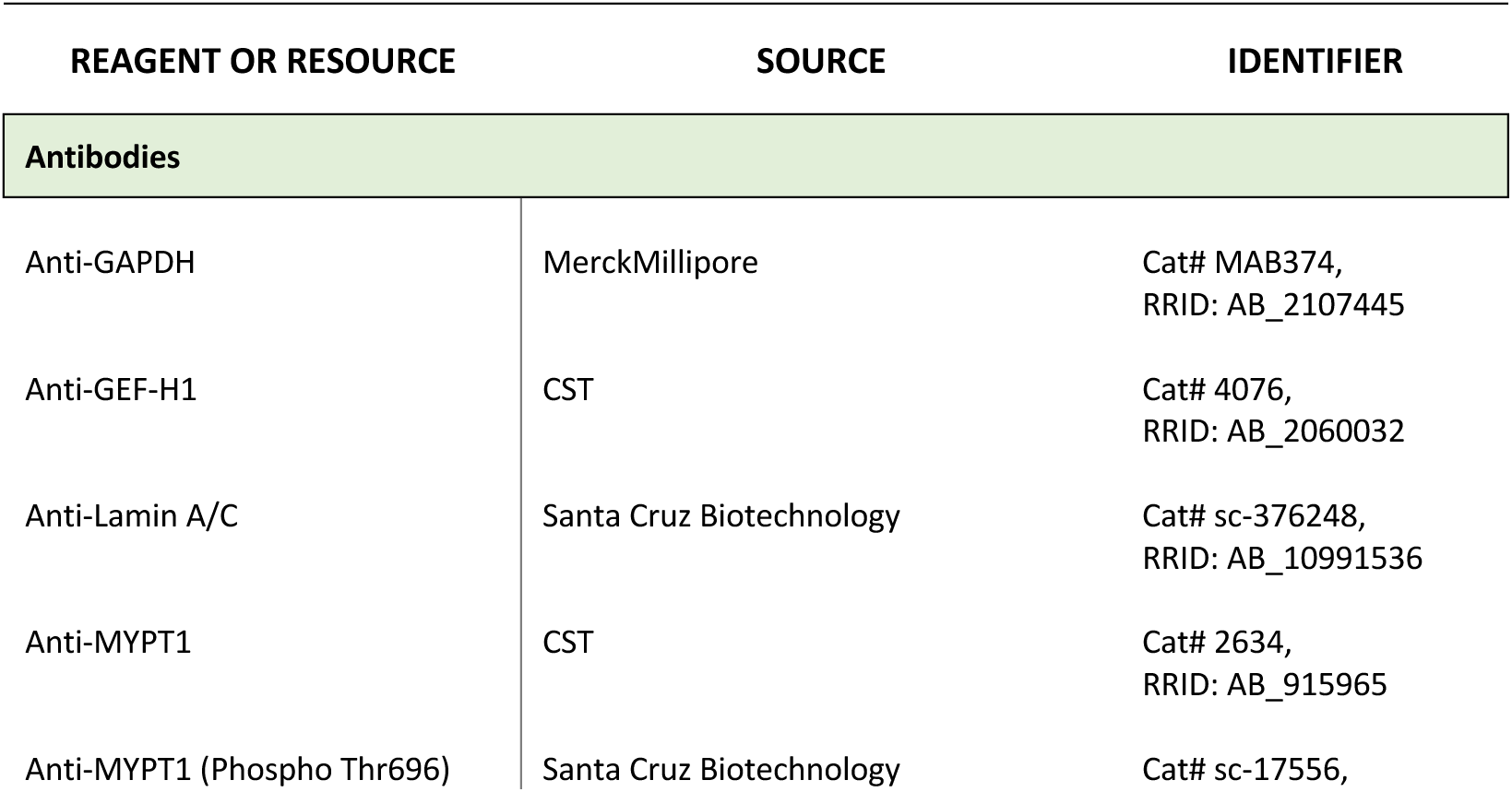
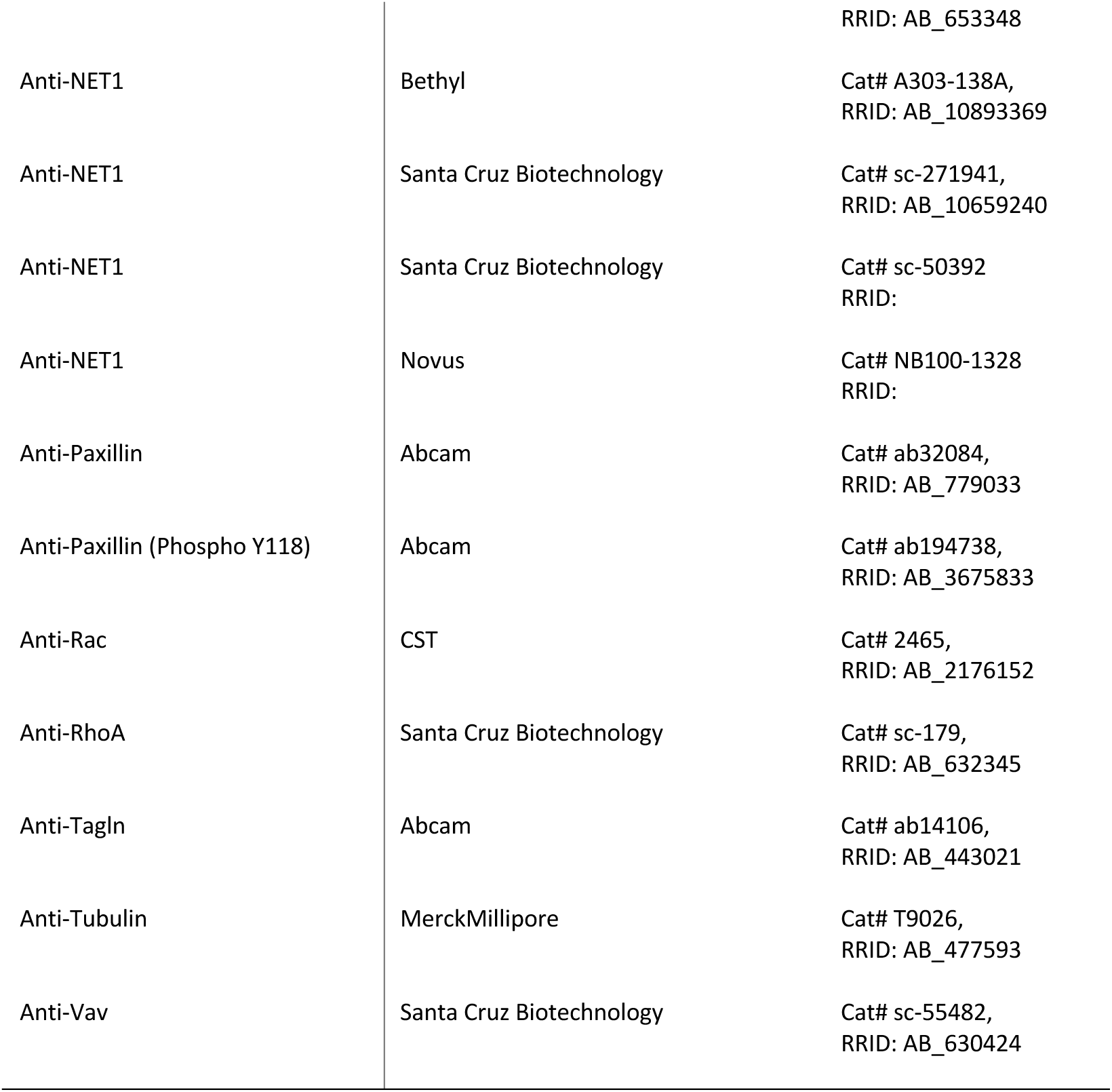
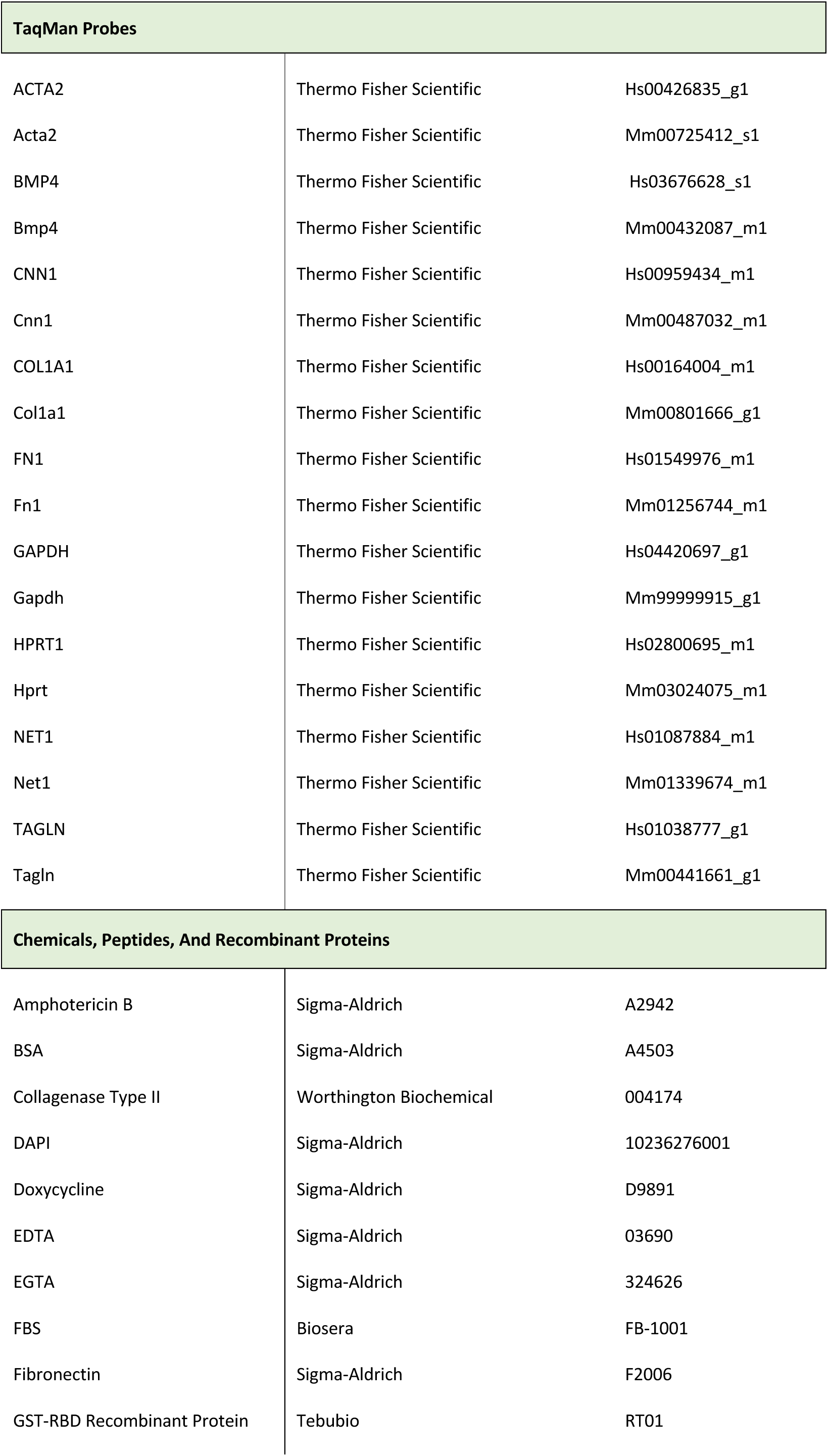
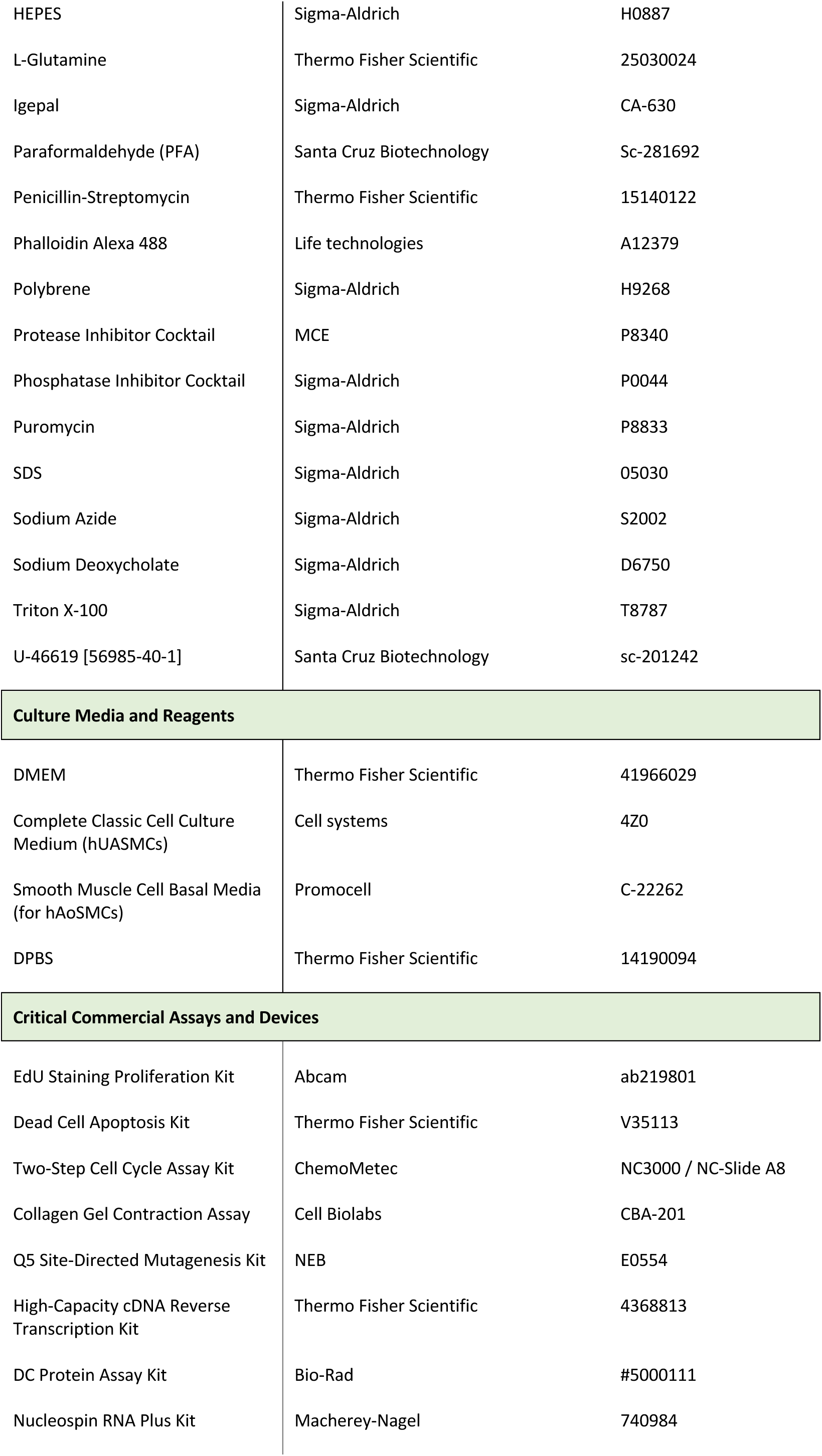
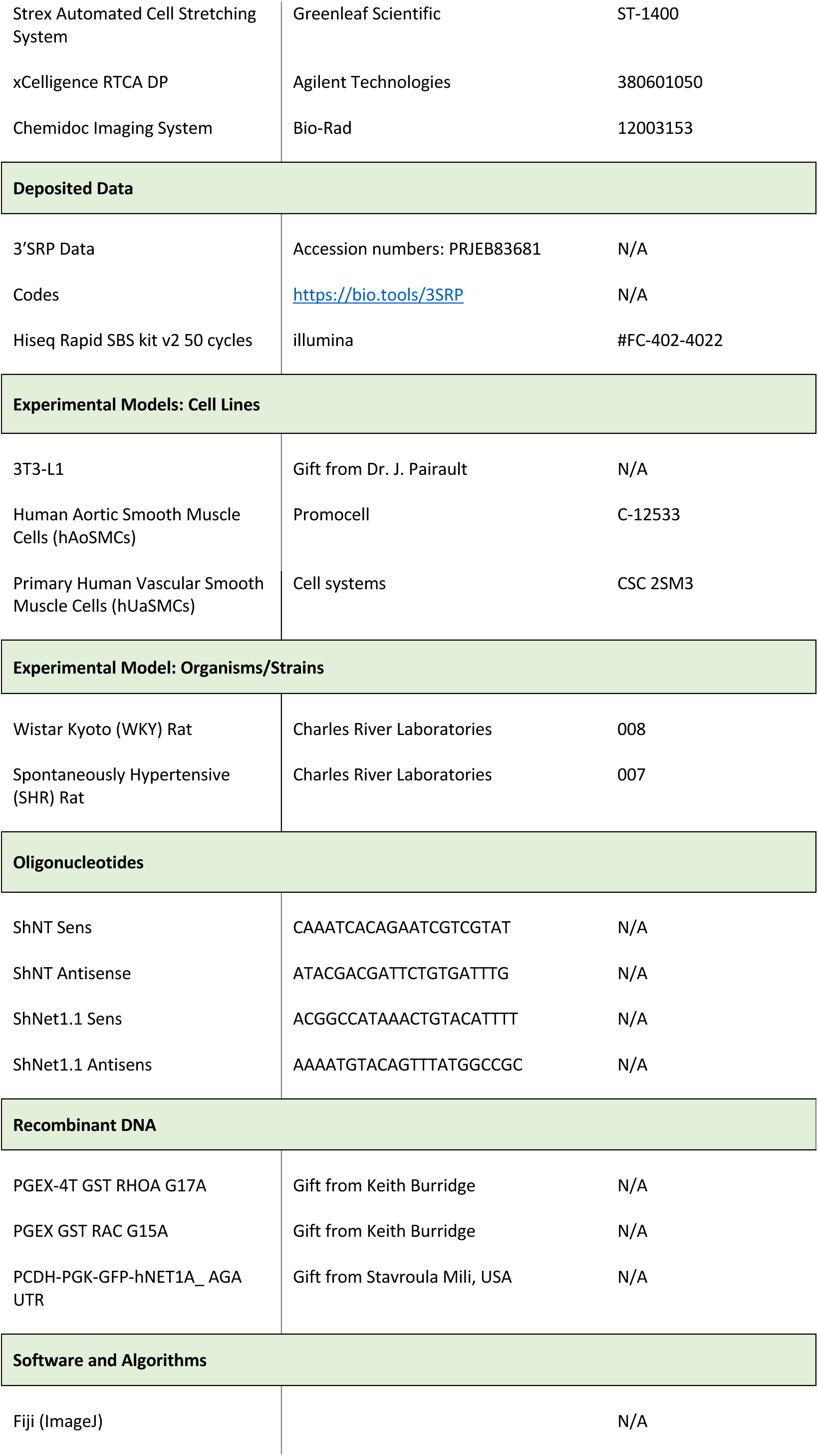
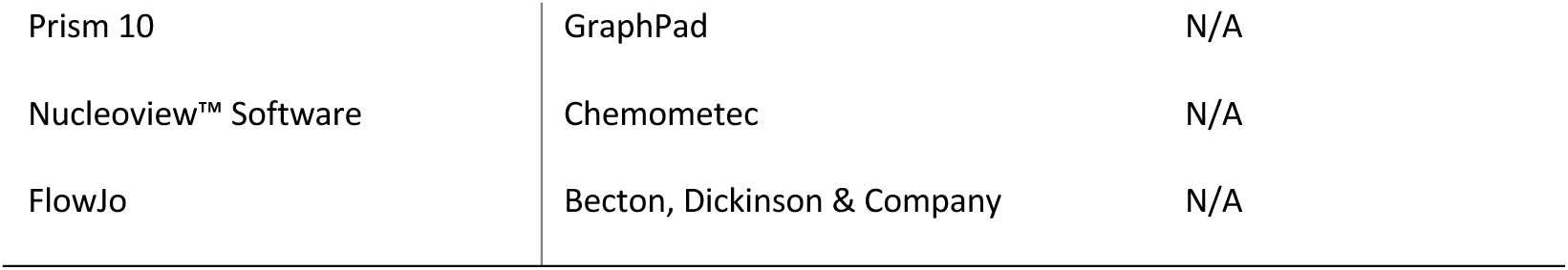

## Lead contact

Requests for further information and resources should be directed to and will be fulfilled by the lead contact, Dr Anne-Clémence Vion, anne-clemence.

## Materials availability

This study did not generate new unique reagents.

## Data and code availability

Data: 3’SRP data will be deposited at the European Nucleotide Archive portal and publicly available in the comming months.

Code: 3’SRP pipeline codes are publicly available at https://bio.tools/ 3SRP as of the date of publication. Accession numbers are listed in the key resources table.

Other items: Any additional information required to reanalyze the data reported in this paper is available from the lead contact upon request.

## ACKNOWLEDGMENTS

We are most grateful to the genomics (GenoA), bioinformatics (BiRD), and microcoscopy (MicroPICell) core facilities (SFR Bonamy, Nantes, France), members of Biogenouest and IBISA, for their expert services. We also thank the Institut Français de Bioinformatique funded by the French National Agency (ANR) (ANR-11-INBS-0013) and the national infrastructure France-Bioimaging (ANR-10-INBS-04), as well as the animal house facility (UTE) of Nantes Université. This work was supported by ANR grants Programme d’Investissements d’Avenir ANR-16-IDEX-0007 (NeXT Initiative), ANR-21-CE14-0016 (to A.-C.V.) and the Lamonica prize (G.L.).

## AUTHOR CONTRIBUTIONS

Conceptualization, A.-C.V.; methodology, A.-C.V.; formal analysis, A.-C.V. and M-A.M.; investigation, M-A.M., M.R., V.V., G.C., E.G., S.P.R.B. and A.-C.V.; data curation, A.-C.V.; writing – original draft, A.-C.V. and M-A.M.; writing – review & editing, A.-C.V., V.S. and G.L.; visualization, A.-C.V. and M-A.M.; supervision, A.-C.V.; funding acquisition, A.-C.V, G.L.

